# The Physiological Response of Salt and Drought-Stressed Plant to Exogenous Application of Salicylic Acid

**DOI:** 10.1101/2022.10.13.512118

**Authors:** Oluwatosin Adebanjo, Elikplim Aku Setordjie, Anelya Almat

## Abstract

Salinity and osmotic stress affect crop growth and yield. To meet the food demand of the increasing global population, there is a need to continually study the plant-stress factor relationship. This experiment studies the physiological response of salt and drought-stressed plant to exogenous application of salicylic acid. Tomato plants were grown in medium, under controlled conditions. The six treatments (T1 - control, T2 - MgSO_4_ for salinity stress, T3 - 5% PEG 8000 for osmotic stress, T4 - control + salicylic acid, T5 - MgSO_4_ + salicylic acid, T6 - 5% PEG 8000 + salicylic acid) were replicated six times to make a total of 36 plants. The treatments were assessed for parameters associated with photosynthetic parameters and yield: chlorophyll content, net photosynthetic rate, leaf water potential, fresh arial weight, leaf ion content, intercellular carbon dioxide concentration, transpiration rate and gaseous exchange. The result showed that the exogenous application of salicylic acid increased the leaf water potential of both the stressed and non-stressed plants. However, for other parameters, the role of MgSO4 and %PEG in inducing salinity stress and drought stress. Respectively, was not clearly observed. Likewise, the exogenous application of salicylic acid showed no clear effect in stressed plants, relative to unstressed plants. Hence, the observations from this experiment showed a high variation in physiological responses and a repeat of the experiment can be considered to further investigate the validation of the role of salicylic acid in plants under salt and osmotic stress conditions.

## INTRODUCTION

Plants are generally exposed to several factors that negatively influence crop production. Some of these factors are biotic and abiotic in nature. Although biotic factors are implicated in global crop production loss (IPPC Secretariat, 2021), abiotic stress factors account for up to 70 per cent loss of staple crops as it presents significant constraints to plant growth and yield (Mantri *et al*. 2012; Basu and Roychoudhury, 2014) and these stress effects have especially been aggravated by the recent menace: climate change (Chaudhry & Sidhu, 2022). A broad array of abiotic stresses, including heat, osmotic stress, cold, and UV can negatively impact the physiological, biochemical, and molecular processes in plants, hence adversely affecting the growth and productivity of plants (Isayenkov, 2012).

Salinity is another abiotic stress factor of global importance due to the increasing salt content in soil on a global scale. Salinity impairs crop growth and development due to its association with osmotic stress. Following an excessive uptake of sodium and chloride ions, salt stress results in the cytotoxicity of plants. In addition to these physiological effects, salinity is typically accompanied by oxidative stress due to the generation of reactive oxygen species (ROS) (Isayenkov, 2012). Generally, the physiological responses of plants to salinity stress have been divided into two main phases. The first is an ion-independent growth reduction, which occurs within minutes to days, causing stomatal closure and inhibition of cell expansion mainly in the shoot (Rajendran et al., 2009). A second phase occurs for a relatively long period, and primarily pertains to the build-up of cytotoxic ion levels, which slows down metabolic processes, reducing leaf water potential, causing premature senescence, and ultimately cell death (Khan *et al*., 2014; Roy *et al*., 2014). Sequel to the reported and the envisaged crop loss resulting from salinity, scientists have researched approaches to mitigating these effects. One of the interesting discoveries is the roles of phytohormones in plant response to abiotic stress conditions, especially salt stress, and osmotic stress. Asides from the common plant hormones - gibberellin, ethylene, auxin and cytokinin, jasmonic acid and salicylic acid have also been studied for their role in plant stress response (Khan *et al*., 2012).

Salicylic acid (SA) is a phenolic compound that is involved in the regulation of the growth and development of plants, and their responses to stress factors. Although SA is commonly implicated in alleviating biotic stress conditions, recent studies have postulated its role in connection with abiotic stress responses (Miuri and Tada, 2014). Miuri and Tada (2014) further reported that SA facilitates the regulation of plant-water relations, photosynthesis, proline metabolism, nitrogen metabolism and antioxidant defence under abiotic stress conditions. The role of SA in strengthening salinity stress-tolerance mechanisms has been extensively evidenced in many crops such as *Vicia faba* (Azooz, 2009), *Brassica juncea* (Nazar *et al*., 2011), *Medicago sativa* (Palma *et al*., 2013), and V. radiata (Khan *et al*., 2014) but needs to be studied more especially on important vegetable crops such as tomato.

Following the predicted continuous increase in salinization and osmotic stress associated to global arable land which is expected to result in 30 per cent land-loss by the end of 2028 (Wang et al., 2003), it is therefore expedient to study the response of plants to salt and osmotic stress, and the role of the exogenous application of salicylic acid on the physiological process of the stressed plants. Hence, this experiment studied the physiological response of salt and osmotic-stressed plant to exogenous application of salicylic acid.

## MATERIALS AND METHODS

### Plant Materials and Growth Conditions

Thirty-six healthy tomato plants (*Solanum lycopersicum*) in 3-4 leaf stage grown in pots filled with moistened coconut fibre. Plants were kindly provided by Polytechnic University Valencia. In order to facilitate the growth and development of a plant’s leafy vegetation, a violet light was installed over the pots.

### Salicylic acid (SA) preparation and application

For stress imposition, the plants were treated with 0.1 mM Salicylic acid by spraying at 7-day intervals. The salicylic acid solution was prepared in water + 0.1% Tween20 and eventually was dissolved (100mM stock) in absolute ethanol. The pots were allocated among six trays (six treatments replicated six times) according to treatments (Table 1).

**Table 1.**
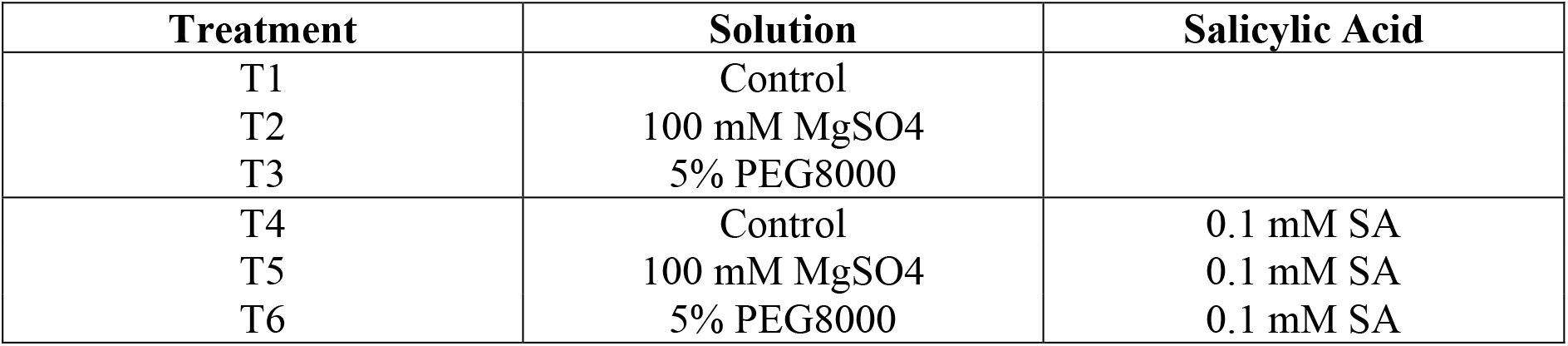
Treatments used in experiment.

### Osmotic stress application

After one week of the experiment tomato plants are subjected to osmotic stress by watering them with a nutrient solution containing polyethylene glycol (PEG 8000) which decrease the water potential. Treated with PEG800 were T3 and T6 samples. Another group (T2 and T5) of plants was treated with magnesium sulphate (MgSO_4_) to cause, in addition to osmotic stress, salt toxicity. Plants sprayed with Hoagland’s solution were serve as a control.

### Water potential

Leaf water potential can be an indicator of plant water stress as a function of soil water availability. The leaf water potential (Ψw) was measured in complete expanded leaflets located in the middle region of the plants. To determine the flow, one leaf per plant was measured using the pressure Scholander chamber techniques.

### Effect on photosynthesis

At 4 weeks, the first photosynthesis parameters measurements were determined. The parameters assessed include biomass fresh weight, chlorophyll content, transpiration weight, intercellular carbon dioxide and gaseous exchange rate of stomata. Other parameters are net photosynthetic rate, ion content in plant, leaf water potential and parameters assessed. Chlorophyll content was established by using SPAD502. In chlorophyll fluorescence, modulated chlorophyll fluorometer (ADC BioScientific Ltd OS5p) was used, being saturation light short pulses at high frequencies (500-30000 Hz). To gas exchange, Infra-red gas analyser (model ADC BioScientific Ltd LCi-SD) was calibrated to supply 0 – 2,400umol of light. A stable, fixed light level can be maintained for all readings, or an automated light response curve can be carried out to characterise a plant’s photosynthetic capacity. All measurements were performed in expanded leaves located in the plant’s middle region.

### Extraction of K^+^, NO_3_^-^ and Mg^2+^

The content of K^+^, NO_3_^-^ and Mg^2+^ was determined with a Spectroquant colorimeter (Merck-Millipore and strips and reagents for Mg2+, NO3 - and K + determinations. Nitrate ions are downgraded to nitrites, which react with an aromatic amine, which reacts with N - (1- naphthyl)-ethylenediamine) to form a red-violet azo dye.

### Data analysis

Shapiro-Wilk test was used to check Normality of data. Due to the inability to attain normality of most parameters, data was submitted to Kruskal-Wallis one way analysis of variance. Agricolae Kruskal package was used to separate means treatments. Data were analysed using R version 4.1.2 and Microsoft Excel.

## RESULTS AND DISCUSSIONS

### Effect of Different Treatments on the Fresh Aerial Weight of Tomato

The different treatments had significant effects on the Fresh weight of above-ground parts (Aerial) of Tomato plants. The highest weight was observed in T1 (70.772 g) followed by T4, T3, T2, T6 and T5 with values of 66.195 g, 50.808 g, 43.747 g, 43.438 g and 42.855 g, respectively (Figure 1).

**Figure 1:**
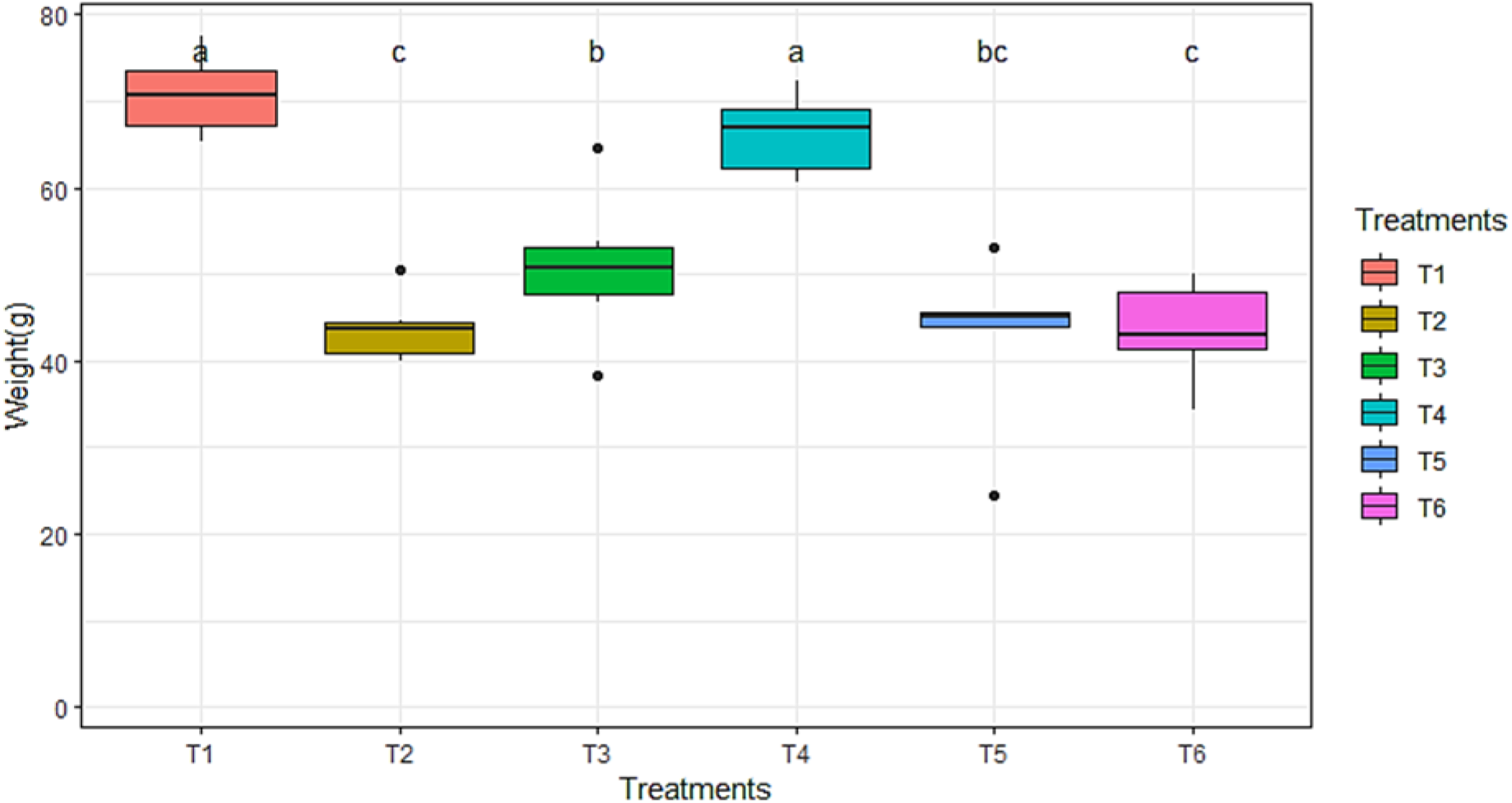
Effect of the various treatments on the Fresh aerial weight of Tomato Plants. *Means followed by the same letters do not differ significantly at p = 0.05

Following previous studies, including that of (Németh *et al*. 2002), an exogenous application of SA at a range of 0.1 - 0.5 mM would increase the tolerance of plants to osmotic stress. However, in the experiment, the SA-treated plants are neutral or reduced in yield in response to the exogenous application of the SA. The reason for this observation is unclear but similar reports have been postulated by Borsani et al. (2001). In his reports, he further opines that the combination of the endogenous SA synthesized by plants in response to defense, together with the exogenous application may potentiate ROS generation in photosynthetic tissues, hence reducing plant performance.

#### Effects of Treatments on Chlorophyll Content of Tomato leaves

Chlorophyll content in the leaves of different treatments of tomato plants were significantly different. The treatments with the highest chlorophyll content were T5 and T3 with SPAD values of 41.1 and 40.567, respectively. These values were followed by those recorded for T2 (36.033), T4 (35.9), T1 (34.933) and T6 (34,533) (Figure 2).

**Figure 2:**
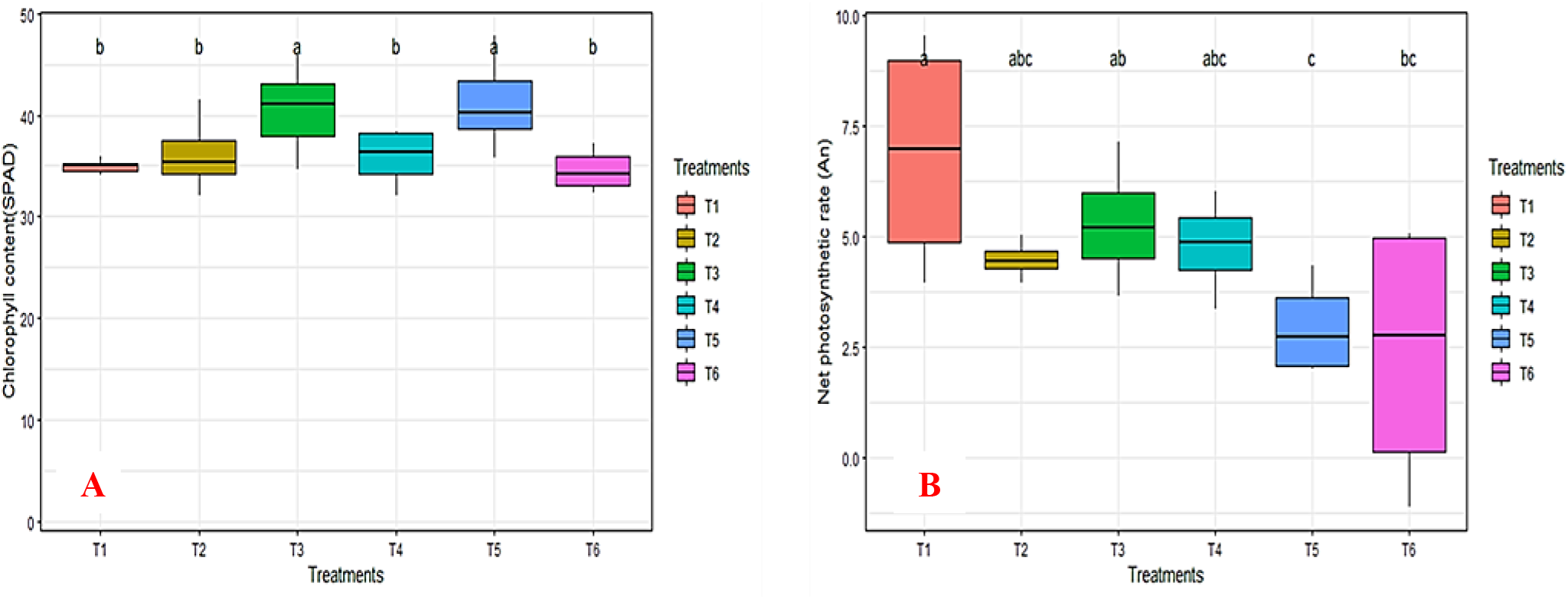
Effect of various treatments on Chlorophyll content(left) and Net photosynthetic rate(right) in Tomato plants. *Means followed by the same letters do not differ significantly at p = 0.05

This agrees with the higher Mg^2+^ content in T5 as magnesium ion is the primary content in chlorophyll (Yan et al., 2001). However, a similar situation was not observed in T2 which recorded a high value for Mg^2+^ content but a relatively lower chlorophyll content.

### Effect of Treatments on Net Photosynthetic Rate of Tomato Plants

There were slight significant differences between the various treatments on tomato plants. T1 had the highest net photosynthetic rate of 6.865 followed by T3, T4, T2, T5 and T6 with values of 5.285, 4.78, 4,47,2.965 and 2.37, respectively, as shown in Figure 2b.

Net photosynthetic rate is adversely affected by water and salinity stress, excess salinity induces osmotic/water stress in plants (Liu, 2011; Liang *et al*., 2019), during such conditions the stomata closes to conserve water reducing the amount of CO_2_ entering leaves thereby reducing plant photosynthesis (Liang *et al*., 2019). This trend is observed in T1 and T3, osmotic stress induced by PEG 8000 (T3) causes Net photosynthetic values to drop slightly in T3. According to Lobato *et al*. (2020), application of salicylic acid increased net photosynthetic rate in stressed tomato plants by reducing the effect of stress on plants and triggering plant defenses, this however is contrary to our results as T4 (Control *Salicylic acid) had a significantly lower net photosynthetic rate when compared to T1 (Control). The lowest Net photosynthetic rate was observed in T6 (PEG 8000* Salicylic acid) indicating that salicylic acid failed to alleviate the effects of water stress induced by PEG. Similarly, the significant reduction in photosynthetic rate in T2 (MgSO_4_) in comparison to T1 shows MgSO4 induced salinity stress in tomato plants, net photosynthetic rate was however not significantly different between T2 and T5 (MgSO_4_*Salicylic acid) indicating the failure of salicylic acid to make tomato plants more tolerant to salinity stress.

### Effect of Treatments on Transpiration rate of Tomato Plants

Transpiration rates recorded in T1 and T4 were significantly higher than the rest of the treatments, with values of 1.45 mmolm^−2^s^−1^ and 1.33 mmolm^−2^s^−1^ respectively. This was followed by T3 (0.745 mmolm^−2^s^−1^), T5 (0.7275 mmolm^−2^s^−1^), T6 (0.6975 mmolm^−2^s^−1^) and T2 (0.535 mmolm^−2^s^−1^) (Figure 3a).

**Figure 3:**
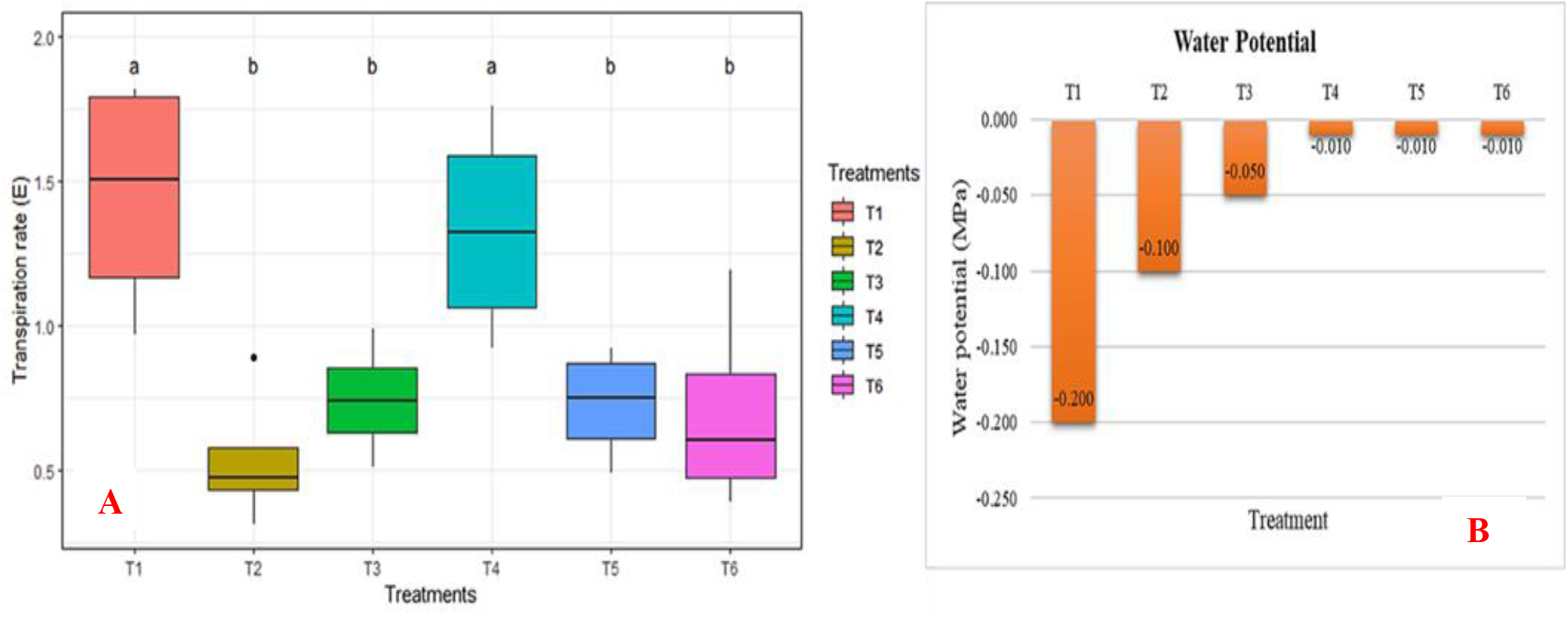
Effect of various treatments on the Transpiration rate(mmolm^−2^s^−1^) (left) and Water potential(right) of Tomato plants. *Means followed by the same letters do not differ significantly at p = 0.05

During water and salinity stress conditions, transpiration rate in palnts in reduced to conserve water in plants. Transpiration rate was significantly lower in all stressed treatments, compared to T1 (Control) and T4 (Control*Salicylic acid). Reduced Transpiration rates in T2 (MgSO_4_) and T5 (MgSO_4_*Salicylic acid) supports reports by Habibi and Sadiqui (2021) which confirms that transpiration in tomato leaves are significantly reduced in by saline conditions due to decreased water absorption by roots. Also, low transpiration rates recorded in T3 (PEG 8000) and T6 (PEG 8000*Salicylic acid) show the negative effect of water stress on tomato plants, in line with Patane *et al*., (2016) which indicated a reduction in transpiration rate of different tomato landraces when there is a water deficit. Low transpiration rates recorded in salicylic acid amended treatments T5 (MgSO_4_*Salicylic acid) and T6 (PEG 8000*Salicylic acid) contradicts reports of salicylic acid reducing effects of water stress and salinity stress in plants (Hayat *et al*., 2020; Naeem *et al*., 2020), and show that application of salicylic acid in water and salinity stress conditions did not improve tolerance of plants to stress.

MPa, higher than values recorded for other treatments. On the other hand, T1, T2 and T3 recorded water potential of −0.200 MPa, −0.100 MPa and −0.050 MPa, respectively (Fig 3b). As supported by Ilyas *et al*., (2017), the higher water potential recorded for T4, T5 and T6 correlated to the exogenous application of SA which placate the osmotic-induced stress in T6, and salt-induced stress in T5.

So, it can be suggested that SA affected the activities of antioxidant enzymes and increased water potential and leaf osmolytes (Askari and Ehsanzadeh, 2015). Hence, the SA helps the plant to manage water potential in osmotic and saline condition.

### Effect of Treatments on gas exchange rate

The results of gas exchange rate (Fig 4a) showed the highest recorded value in T4 and this is not significantly different from value recorded in T1. T2 recorded the lowest gas exchange rate but showed no significant different from value recorded in T5 and T6. Additionally, T3 recorded a value higher that those recorded in T5 and T6, but it is not significantly different from T5 and T6. According to Stevens *et al*., (2006) and several other research, the application of SA to salt-stressed and osmostic-stressed plants is expected to cause am increase in leaf gas exchange rate. However, this experiment presents a different result but the reason for this anomaly is not yet clear. The net photosynthetic rate is also expected to be positively correlated to gaseous exchange and intercellular carbondioxide concentration (Khan *et al*., 2014), but this is not the case in this experiement.

**Figure 4:**
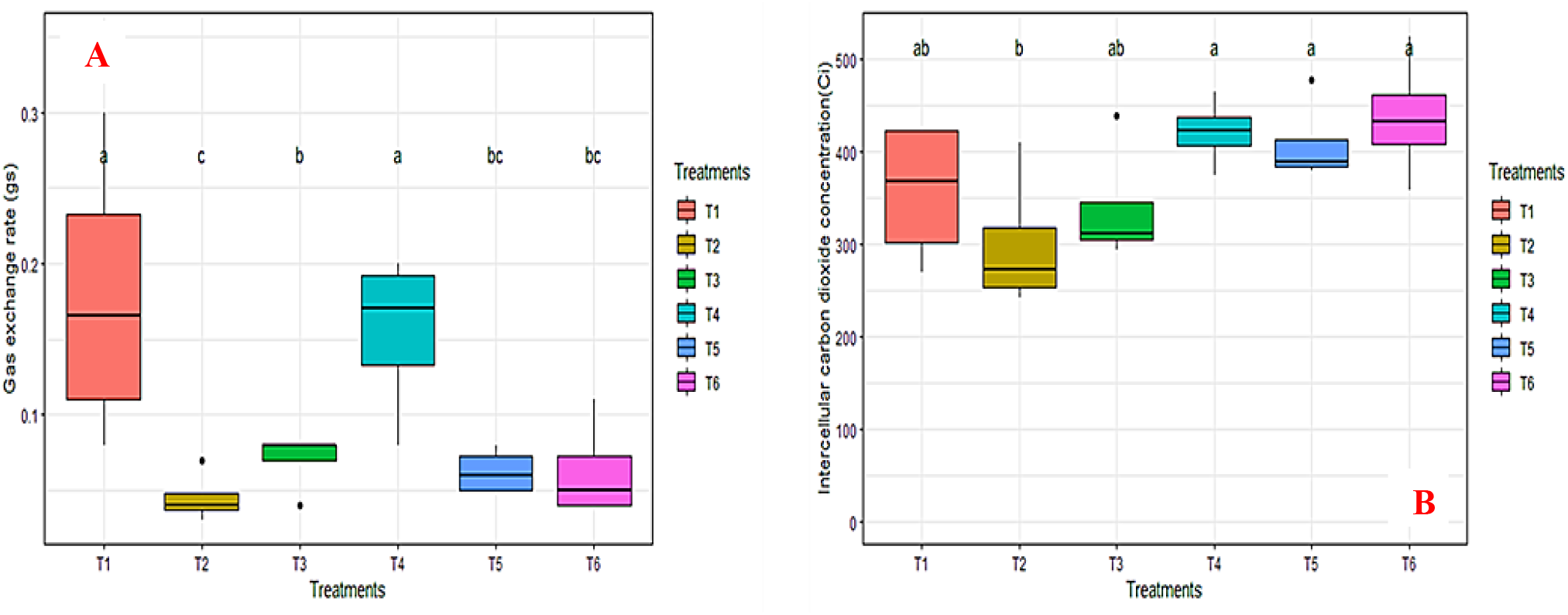
Effect of various treatments on Gas exchange rate(mmol^−2^s^−1^) (left) and Intercellular carbon dioxide(right) in Tomato plants. *Means followed by the same letters do not differ significantly at p = 0.05

### Effect of Treatments on intracellular carbon

For intracellular carbon dioxide (Fig 4b), T6 recorded the highest value with no significant difference from T1, T3, T4, and T5. However it is significantly different from T2 which recorded the lowest value for intercellular carbon dioxide. Generally, the plants treated with SA have higher intercellular carbon dioxide concentrations compared to those without the application of SA. This observation is in tandem with the report of Khan *et al*., (2014) where they reported that stressed plants, especially salt-associated stress, can influence a reduction of 23 - 30% in the intercellular carbon dioxide concentration. This is particularly clear in a comparison between plants with SA and plants without SA application as in the case between T1 and T4, T2 and T5, and between T3 and T6.

### Effect of various Treatments on the Ion content

The different treatments had no significant effects on the ion content of the plants (Table 3). The non-stressed plants, T1 had the highest NO_3_^-^ content of 76.5 mg, followed by T4 with a NO_3_^-^ content of 43.5 mg. While T1 and T4 had higher NO_3_^-^ content, they are not significantly different from other treatments. Relative to T1 and T4, the reduced but insignificantly different reduction on NO_3_^-^ corroborate the report from Meloni *et al*., (2014). For the magnesium contents, T2 and T5 had the best performance, and this is positively correlated with the MgSO_4_ added to the T2 and T5 to induce salinity stress. In comparison between the stressed and the unstressed plants, there is no significant difference in the K+ content recorded. However, a trend of seemingly elevated K^+^ under Mg reduction was observed and this corroborates the work of Jezek and Geilfus (2015).

**Table 3:** Effect of various treatments on the Ion content of Tomato plants.

| **Treatment** | **$NO_3$ (mg)** | **$Mg^{2+}$ (mg)** | **$K^+$ (g)** |
| --- | --- | --- | --- |
| T1 | 76.5<sup>a</sup> | 30.5<sup>ab</sup> | 0.475<sup>ab</sup> |
| T2 | 25.5<sup>a</sup> | 83.5<sup>a</sup> | 0.39<sup>c</sup> |
| T3 | 15<sup>a</sup> | 31<sup>ab</sup> | 0.51<sup>a</sup> |
| T4 | 43.5<sup>a</sup> | 29<sup>b</sup> | 0.505<sup>ab</sup> |
| T5 | 16.5<sup>a</sup> | 78<sup>a</sup> | 0.445<sup>bc</sup> |
| T6 | 12<sup>a</sup> | 28<sup>b</sup> | 0.46<sup>abc</sup> |

\*Means followed by the same letters do not differ significantly at $p = 0.05$

## CONCLUSION

Following the results obtained in the experiment, the application of exogenous SA improves the water potential of stresses plant (salinity and osmotic stress). However, the effect of the exogenous application of salicylic acid on other physiological processes of salt-stress and osmotic-stressed tomato plant is not clear. The physiological responses of the plants vary grateful to salt stress, osmotic stress, and exogenous application of salicylic acid. Although other researchers postulated that the application of salicylic acid to salt-stressed, and osmotic stressed plants attenuate the abiotic stress effect, this experiment recorded irregular responses, neither positive nor negative. The mechanism for this anomaly response is not clearly identified yet, hence, the experiment could be repeated to confirm is there is consistency in the neutral response.

## Supporting information

Supplementary raw data

